# Effect of egg cannibalism on mating preferences and reproductive fitness of *Menochilus sexmaculatus* Fabricius (Coleoptera: Coccinellidae)

**DOI:** 10.1101/2020.03.14.991695

**Authors:** Tripti Yadav, Omkar, Geetanjali Mishra

## Abstract

Cannibalism has been reported in a large proportion of coccinellids in fields as well as in laboratories but studies involving mate preferences and potential benefits of cannibalism on reproduction in *Menochilus sexmaculatus* Fabricius (Coleoptera: Coccinellidae)have yet not been done. Thus, we assessed the effect of conspecific egg cannibalism on mate preferences and reproductive outputs including offspring development. Higher mate preferences were recorded for non-cannibal mates (fed on *A. craccivora*) than cannibal ones (fed on conspecific eggs). Mating parameters significantly influenced by cannibalism. Time to commence mating lasted less for homogeneous diet pairs than heterogeneous diet pairs. Longer copulation duration and higher fecundity were recorded when one of the individuals in mating pair or both was a non-cannibal. Egg viability did not differed significantly in all reciprocal crosses. Total developmental durations of offspring were similar for all mating pairs.

## Introduction

Mate preferences are often plastic in nature and they change in response to changing environment (Polis, 1981; Candolin et al., 2016; Rosenthal, 2017) and internal conditions of the mates (Srivastava and Omkar, 2004; Perry et al., 2009; Omkar and Singh, 2010). An individual’s mate preference is thus a result of a combination of multiple traits including the quality of their potential mate (Hebets et al., 2008; Schultzhaus et al., 2017). Quality of the individual is known to be affected by multiple factors such as age (Avent et al., 2008; Anjos-Duarte et al., 2011), size (Downhower and Brown, 1980; Tina and Muramatsu, 2015; Locatello et al., 2016), immunity (Landis et al., 2015; Iglesias-Carrasco et al., 2017), sperm quality and quantity (Pizzari and Birkhead, 2002; Jones and Elgar, 2004). Of these, the latter three are especially known to be modulated by diet (Lewis and Wedell, 2007; Chang et al., 2008). In addition, in insects, the reproductive value of potential mates is assessed by their chemical signatures (Johansson and Jones, 2007; Peterson et al., 2007), which also tend to change with diet (Fedina et al., 2012, Otte et al., 2015). Thus, dietary history of an individual is likely to change their preferability as a mate or their competitive performance during mate selection. In nature, access to good quality food is often limited and there is extensive competition for it (Prokopy et al., 1984; Polis and Holt, 1992; Yasuda et al., 2001). In individuals with cannibalistic tendencies, limitation on good quality resource may also lead to increased rate of intraspecific predation. Intraspecific predation is also known to serve as a strategy for reproductive competition by reducing the fitness of other individuals of the same sex, directly by cannibalizing sexual competitors or indirectly by eating their offspring (Kynard, 1978; Ito et al., 2000) and cannibalizing potential mates (Elgar, 1992; Lawrence, 1992; Wu et al., 2013; Schutz and Taborsky, 2005).

Cannibalism though a known means of reproductive competition is also known to negatively affect the fitness of individuals produced (Dimetry, 1974; Chapman et al., 1999). On the other hand, it has also been suggested that the natural selection might favour cannibals over non-cannibals as the studies in spider species have shown that cannibalism can have an indirect effect on their fitness by boosting their fecundity and offspring viability (Wu et al., 2013; Pruitt et al., 2014). Studies in *Tribolium confusum* has also revealed that cannibalism can be inherited as a trait (Steven, 1994). While cannibalism is largely believed to occur in unsuitable abiotic and biotic environments (Agarwala and Dixon, 1992; Wagner et al., 1999; Svenning and Borgstrom, 2005; Rojht et al., 2009), multiple studies indicate an increased expression of cannibalism even under suitable conditions (Steven and Mertz, 1985; Smith and Reay, 1991; Hecht and Pienaar, 1993; Stevens, 1994; Baur, 1994; Rudolf et al., 2010).

In view of the largely believed lower fitness levels of cannibals, and also the inheritance of cannibalistic traits, we though it worth investigating whether a cannibalistic dietary history would modulate the mate preference across sexes. For this, we conducted studies on an aphidophagous ladybird beetle, *Menochilus sexmaculatus* (Fabricius), which is highly cannibalistic in nature. Also, multiple studies on pre- and post-copulatory sexual selection have been conducted on this ladybird. Ladybird beetles are known to show non-random mate preferences based on morph (Saeki et al., 2005; Srivastava and Omkar, 2005; Bista and Omkar, 2015), size (Brown, 1990a, b), food conditions (Ueno et al., 1998; Dubey, 2016) and age (Pervez et al., 2004; Srivastava and Omkar, 2004; Omkar and Singh, 2010).

It is our hypothesis that cannibals should not be preferred in general by mates having a non-cannibalistic dietary history. However, it is possible in view of the inheritability of cannibalistic traits and also likely similarity in cuticular hydrocarbon profiles of cannibals owing to similar diets, that cannibalistic mates might prefer cannibals as mates. We also propose that adults with similar dietary histories are likely to have offspring with higher fitness levels than those with different dietary histories.

## Materials and Methods

### Stock culture

Adults of *Menochilus sexmaculatus* (*n*=40) were collected from the local agricultural fields of Lucknow, India (26°50’N, 80°54’E). This species was selected as an experimental model due to its abundance in local fields, wide prey range and high reproductive output (Omkar and Bind, 2004; Omkar et al., 2005). Adults were fed with *ad libitum* supply of cowpea aphid, *Aphis craccivora* Koch (Hemiptera: Aphididae). This aphid was established on *Vigna unguiculata* L. plants in the glasshouse (25 ± 2°C, 65 ± 5% R H). Adults were paired and placed in Petri dishes (hereafter, 9.0 × 2.0 cm), which were kept in Biochemical Oxygen Demand incubators (Yorco Super Deluxe, YSI-440, New Delhi, India) at 25 ± 1°C, 65 ± 5% R.H., 14L: 10D. Eggs laid were collected, and held in new plastic Petri dishes until hatching, which usually occurs within 2-3 days. Once the first instars began moving on or away from the remnants of their egg clutch, they were gently removed using a fine camel-hair paintbrush and assigned individually to clean experimental Petri dishes (size as above).

### Cannibalism and mate preference

To assess the effect of cannibalism on mating success of adults, first instar larvae were collected from stock culture and randomly assigned for rearing till pupation on either one of two different dietary regimes, (i) conspecific eggs, (Yadav et al., 2019) and (ii) aphids (*ad libitum*, 40 mg). Adults that emerged from these larvae were sexed, paired and placed on the same regimes as their immature stages. Offspring of these adults were reared on their parental diet regime and then subjected to mate preference trials at the age of ten days.

For male mate preference trials, a ten-day-old unmated male reared on conspecific eggs was provided with a simultaneous choice of two virgin females of same age, one from each dietary regime (*i.e.* conspecific eggs and aphids). To differentiate females of above food regimes they were colour marked with small dots of non-toxic green or blue colour on the posterior edge of their left elytra (Dubey, 2016). To prevent any bias in mate preference due to these colours, the marking was switched in each replicate. If mating occurred, the rejected females were removed and mating was allowed to complete. The same procedure was followed in case of aphid diet. In case of female mate preference trials, females of each dietary regime were provided simultaneous choice of males from both dietary regimes. Each mate preference treatment was replicated 20 times.

The mating parameters, *viz.* time of commencement of mating and copulation durations of each pair were recorded. In order to assess the effect of cannibalism on reproductive performance, the following combinations obtained from both the female and male mate preference setup were observed, F_nc_ × M_nc_, F_nc_ × M_c_, F_c_ × M_nc,_ F_c_ × M_c_ (F and M representing female and male respectively; ‘nc’ and ‘c’ representing non-cannibal and cannibal, respectively).

After the termination of mating, the male was removed and the females were maintained on their pre-assigned diet. Daily oviposition and hatching were recorded for the next 5 days

### Statistical analysis

The chi-square (*χ*2) goodness-of-fit analysis was used to test the null hypothesis of random mating. Data on mating and reproductive outputs, i.e. time of commencement of mating, copulation duration, fecundity, egg viability, and developmental durations were first tested for normality (Kolmogorov-Smirnoff). On being found normally distributed, each of the previous measurements was subjected to one-way analysis of variance (ANOVA) with diet as independent factors. The analysis was followed by the comparison of means using post hoc Tukey’s test at 5% probability level. All statistical analyses were conducted using R 3.6.1. statistical software.

## Results

Non-cannibal adults were preferred significantly as mates by non-cannibals (F_nc_: χ^2^= 5, df= 1, P<0.05; M_nc_: χ^2^=7.2, df= 1, P<0.05). However, cannibals did not showed any preference (F_c_: χ^2^= 1.8, df=1, P>0.05; M_c_: χ^2^=0.8, df= 1, P>0.05) (Table 1) for cannibals or non-cannibals.

**Table 1.**
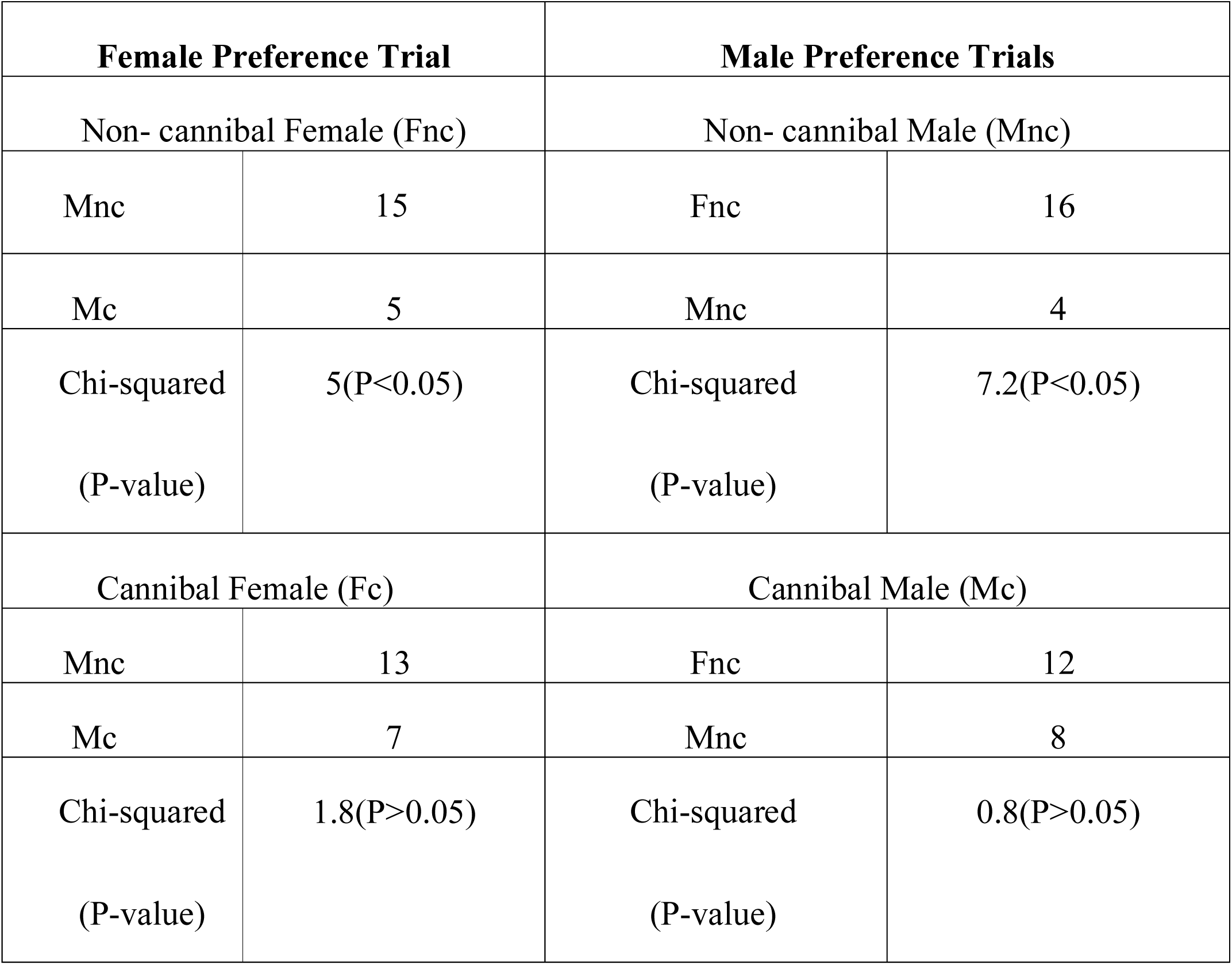
Effect of egg cannibalism on mate preferences of female and male *Menochilus sexmaculatus.* † ‘F’ and ‘M’ representing female and male respectively; ‘nc’ and ‘c’ representing aphid and conspecific egg diet, respectively.

| **Female Preference Trial** |  | **Male Preference Trials** |  |
| --- | --- | --- | --- |
| Non- cannibal Female (Fnc) |  | Non- cannibal Male (Mnc) |  |
| Mnc | 15 | Fnc | 16 |
| Mc | 5 | Mnc | 4 |
| Chi-squared<br>(P-value) | 5(P<0.05) | Chi-squared<br>(P-value) | 7.2(P<0.05) |
| Cannibal Female (Fc) |  | Cannibal Male (Mc) |  |
| Mnc | 13 | Fnc | 12 |
| Mc | 7 | Mnc | 8 |
| Chi-squared<br>(P-value) | 1.8(P>0.05) | Chi-squared<br>(P-value) | 0.8(P>0.05) |

Time to commence mating (TCM) was significantly influenced by diet regime of mates in both female (F_3,56_=8.729, P<0.05) and male (F_3,56_=4.452, P<0.05) preference trials with decreased TCM for homogenous diet pairs and increased TCM for heterogeneous diet pairs regardless the adults were cannibal or non-cannibal pairs (Figure 1).

**Figure 1.**
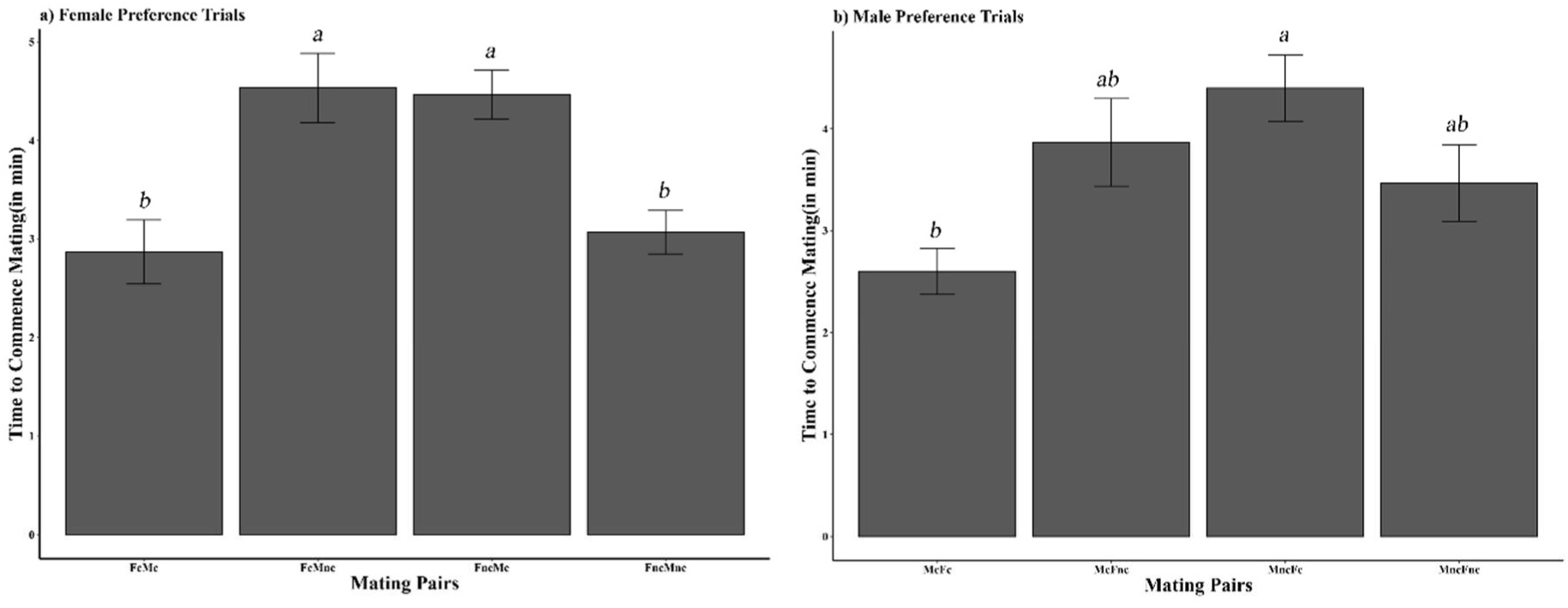
Time to commencement of mating in a) female and b) male mate preference trials when male and female mated with a cannibal and non-cannibal mate in *Menochilus sexmaculatus.* Values are mean ± SE. Similar letters indicate lack of significant difference (P-value > 0.05). † ‘F’ and ‘M’ representing female and male respectively; ‘nc’ and ‘c’ representing aphid and conspecific egg diet, respectively.

Copulation duration was also significantly influenced by mate diet regime in both female (F_3,56_= 12.98, P<0.05) and male (F_3,56_=8.55, P<0.05) preference trials. Shortest copulation duration in both female and male preference trials was recorded for heterogeneous diet pairs whereas longest duration was recorded for non-cannibal pairs (Figure 2).

**Figure 2.**
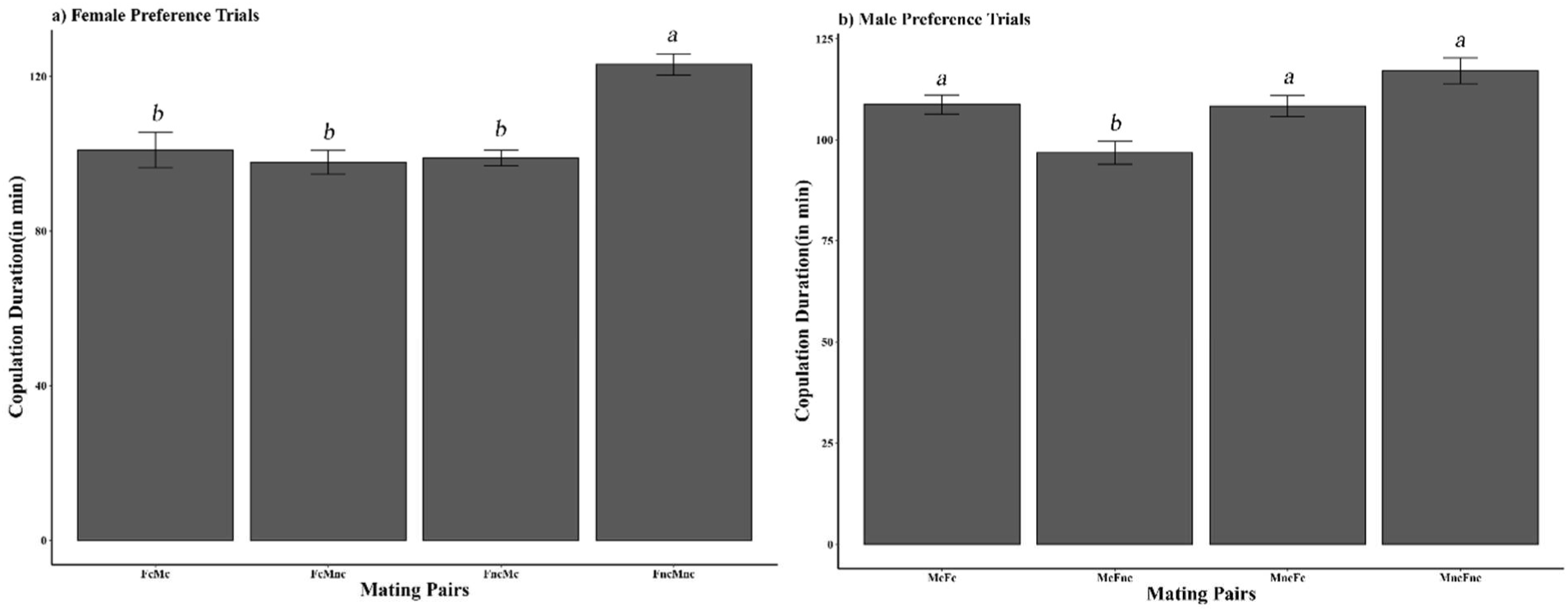
Copulation duration in a) female and b) male mate preference trials when male and female mated with a cannibal and non-cannibal mate in *Menochilus sexmaculatus.* Values are mean ± SE. Similar letters indicate lack of significant difference (P-value > 0.05).

Fecundity was significantly influenced by diet regimes of mates in both female (F_3,56_=120.5, P<0.05) and male (F_3,56_=136.2, P<0.05) mate preference trials. Non-cannibal females were more fecund than cannibals were; pairing with non-cannibal males also boosted egg production (Figure 3).

**Figure 3.**
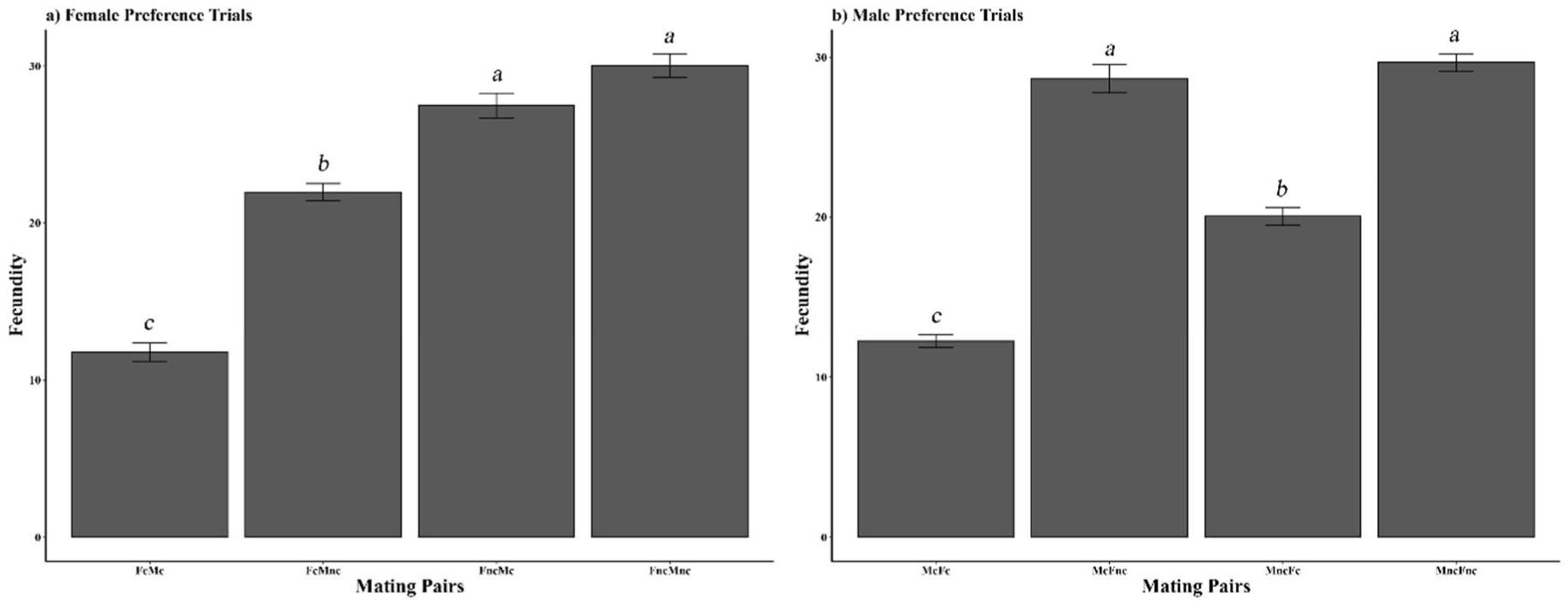
Average fecundity in a) female and b) male mate preference trials when male and female mated with cannibal and non-cannibal mate in *Menochilus sexmaculatus.* Values are mean ± SE. Similar letters indicate lack of significant difference (P-value > 0.05).

Percent egg viability was not significantly influenced by adult diet in both female (F_3,56_=0.26 P>0.05) and male (F_3,56_=0.42 P>0.05) (Figure 4) mate preference trials.

**Figure 4.**
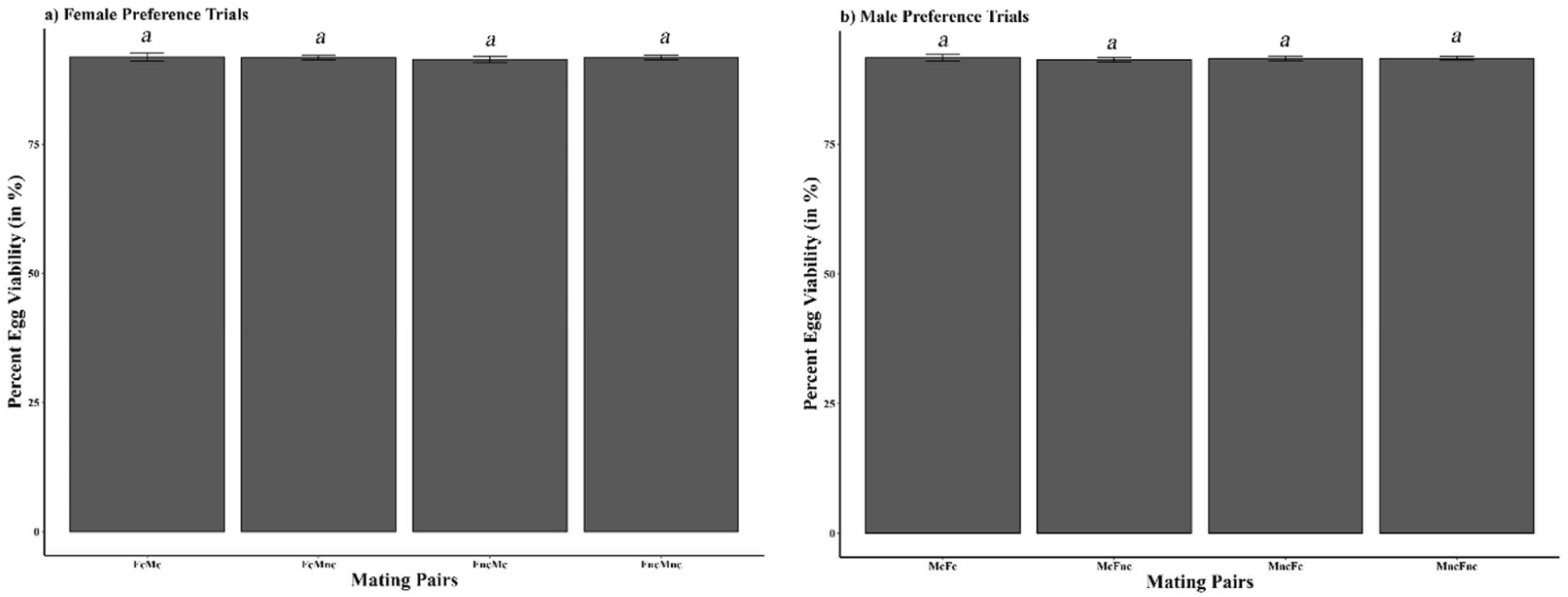
Percent egg viability in a) female and b) male mate preference trials when male and female mated with a cannibal and non-cannibal mate in *Menochilus sexmaculatus.* Values are mean ± SE. Similar letters indicate lack of significant difference (P-value > 0.05).

Total developmental duration of offspring was found similar in all mating treatments regardless of parents’ diet regime (F_3,76_=2.566, P>0.05) (Figure 5).

**Figure 5.**
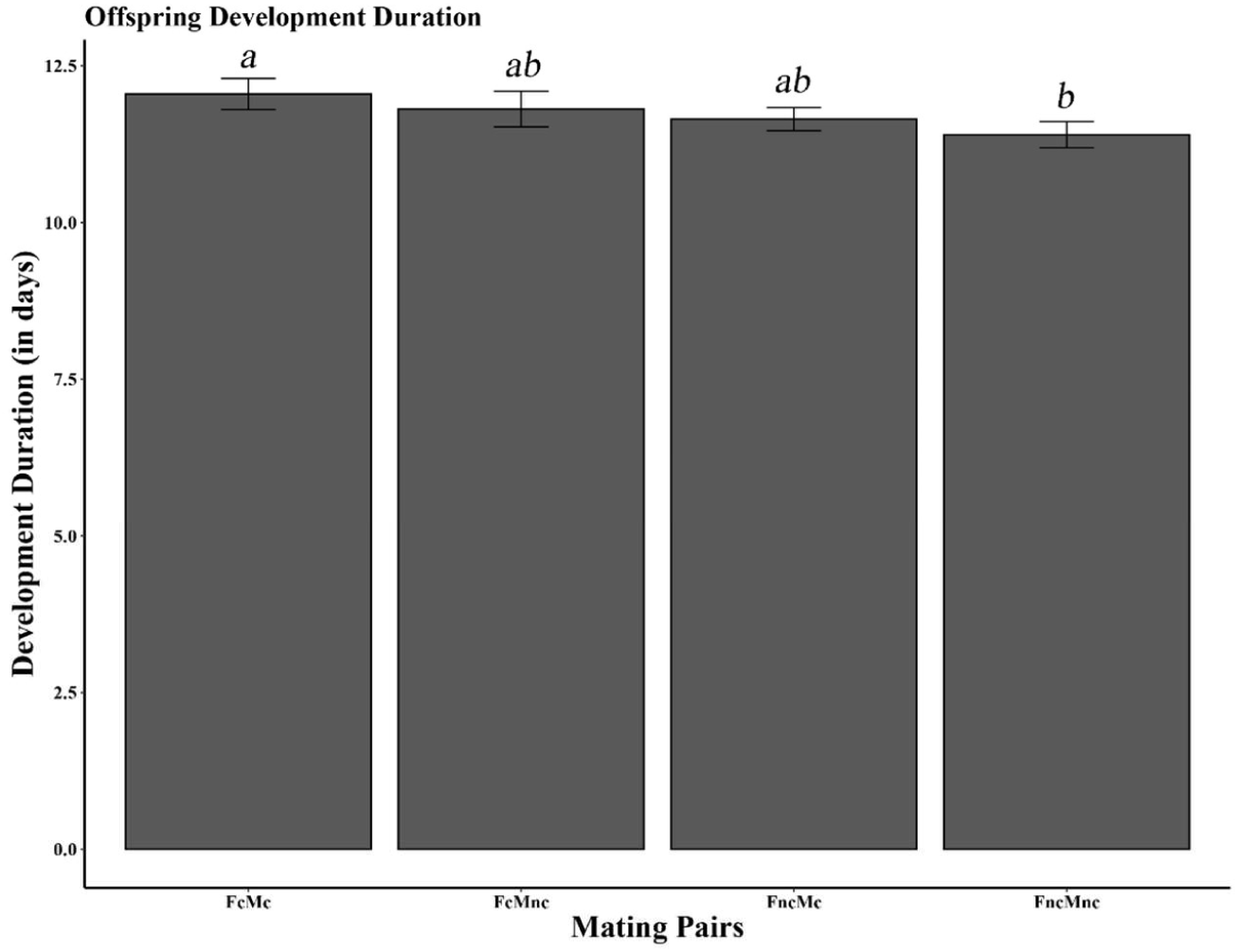
Effect of parental egg cannibalism and mating structure on total developmental durations. Values are mean ± SE. Similar letters indicate a lack of significant difference (P-value > 0.05).

## Discussion

The present study revealed that non-cannibal adults of *M. sexmaculatus* preferred non-cannibal mates and this preference was evident in the mating parameters. The reproductive output was also affected by parental diet and was supportive for the preference of non-cannibals as mates. However, total developmental durations of offspring were found to be similar for all mating pairs.

Higher mating preferences recorded for non-cannibal adults of *M. sexmaculatus* in both female and male preference trials can be attributed to their higher fitness levels owing to their better growth, development and reproduction (Agarwala and Yasuda, 2000; Omkar and Bind, 2004) on aphids, in particular, *A. craccivora*, which is the most preferred prey (Omkar and Bind, 2004). Individuals across taxa are known to pursue the mate of the highest quality in order to increase the quantity and quality of offspring. Another, possible reason of avoiding cannibal mates may be attributed to greater propensity of cannibals to cannibalize their own offspring (Smith and Reay, 1991; Hecht and Pienaar, 1993) which can be detrimental as it can result in decreased fitness (Hamilton, 1964a, b; Elgar and Crespi, 1992; Joseph et al., 1999). Wade (1980) reported that the cannibalistic trait in flour beetle, *Tribolium confusum* du Val was significantly influenced by mating structure, *i.e.* within-group and random mating, with increased rate of cannibalism within-group mating treatments.

Longer and shorter TCM for heterogeneous and homogenous diet pairs respectively, in this case, may be a result of cuticular hydrocarbon-mediated mating signals. Cuticular hydrocarbon (CHCs) plays a major role in kin and mate recognition (Geiselhardt et al., 2009, 2012) in insects and both environmental and internal factors are known to modulate the CHCs especially diet (Brazner and Etges, 1993; Etges, 1992; Etges et al., 2006). Studies in *Drosophila melanogaster* (Sharon et al., 2010) have reported that the commensal bacterial associated with the food media resulted in altered CHCs profiles of the adult flies and resulted in diet-specific assortative mating.

Shorter copulation duration was recorded for homogenous cannibal pairs as compared to homogenous non-cannibal or heterogeneous pairs. Mating is an energy-intensive process, can probably not be sustained long enough by cannibal adults, owing to relatively poor nutritive profile of eggs compared to aphids, leading to early mating termination. Another possible reason can be that cannibal adults are avoided as mates, as it may result in offspring with high cannibalistic tendencies, thereby causing increased probability of future loss of inclusive fitness (Hamilton, 1964a, b; Joseph et al., 1999). Stevens (1994) reported that the cannibalistic trait can be inherited in a Mendelian genetic pattern and can increase the expression of cannibalism in *T. confusum* population.

Non-cannibal females were more fecund than cannibal ones, with non-cannibal males further enhancing egg production. The high fecundity of non-cannibal females is more likely because of the nutritional suitability of aphids (Omkar and Bind, 2004; Omkar et al., 2005; Srikanth and Lakkundi, 1990; Agarwala and Yasuda, 2000) and the efficient utilization and conversion of aphid biomass rather than conspecific egg biomass into egg (Ware et al., 2008). The role of the non-cannibal male in enhancing egg production suggests that males do contribute to egg production which was earlier believed to be a female domain (Lewis et al., 2011; Michaud et al., 2013). It is likely that nutrition might enhance the quality as well as quantity of seminal fluid proteins (Perry and Rowe, 2010; Michaud et al., 2013) resulting in higher egg production. A similar pattern of fecundity was observed in *Adalia bipunctata* (Linnaeus) (Dimetry, 1974). However, in *Coccinella undecimpunctata* L. while the lowest fecundity was observed in cannibal mating pair as in this study, highest fecundity was obtained when males cannibalized eggs (Bayoumy et al., 2016). Studies in *Hippodamia convergens* Guérin-Méneville (Bayoumy and Michaud, 2015) reported greater reproductive benefits of cannibalism with cannibals being more fecund than non-cannibals.

Percent egg viability was found to be similar in all mating treatments, which is not on the lines of our expectations of decreased performance in cannibals. In contrast, studies in *H. convergens* (Bayoumy and Michaud, 2015) and *C. undecimpunctata* (Bayoumy et al., 2016) reported higher percent egg viability for cannibal pairs as compared to non-cannibal ones. The increase in percent egg viability in these studies may be attributed to diet complementation as the cannibal pairs were provided with mixed diet *i.e. Ephestia kuehniella* and conspecific eggs. The insignificant difference in total development duration of offspring in all the mating combinations suggests efficient utilization of both aphid and conspecific egg by larval instars to complete their development. A similar study in *Coccinella undecimpunctata* L. (Bayoumy et al., 2016) reported shorter offspring developmental duration (by accelerating pupation) when both the parents cannibalized eggs. Earlier studies in coccinellids have shown stage-specific (Bayoumy and Michaud, 2015a; Bayoumy et al., 2016; Omkar et al., 2004; Kumar et al., 2014) benefit of cannibalism. In *Coleomegilla maculata* Lengi (Gagne et al., 2002) cannibalism by second instar larvae resulted in shorter development duration and better larval growth than non-cannibal.

In conclusion, our result supports the hypothesis that cannibals should not be preferred by mates having non-cannibalistic dietary history and that the non-cannibals were better performers in terms of mating and reproductive parameters. However, the parental diet had no effect on the development of the offspring. Conspecific eggs in spite of being highly nutritive resulted in poor performance of cannibals. This was contradictory to previous studies where cannibalism enhanced the performance of adults and their offspring. Cannibalism appears to be an adaptive strategy for nutrient imbalance and seems to be detrimental by lowering the selective value of cannibals.

## Acknowledgements

TY gratefully acknowledges CSIR, New Delhi, India, for Junior Research Fellowship, (09/107(0405)/2019-EMR-I) dated July 01, 2018.

